# The genome and mRNA transcriptome of the cosmopolitan calanoid copepod *Acartia tonsa* Dana improve the understanding of copepod genome size evolution

**DOI:** 10.1101/560102

**Authors:** Tue Sparholt Jørgensen, Bent Petersen, H. Cecilie B. Petersen, Patrick Denis Browne, Stefan Prost, Jonathon H. Stillman, Lars Hestbjerg Hansen, Benni Winding Hansen

**Author notes:** Lars Hestbjerg Hansen, Department of Environmental Science - Environmental Microbiology and Biotechnology, Aarhus University, Roskilde, Denmark, Tue Sparholt Jørgensen Department of Science and Environment, Roskilde University, Roskilde, Denmark.

## Abstract

Members of the crustacean subclass Copepoda are likely the most abundant metazoans worldwide. Pelagic marine species are critical in converting planktonic microalgae to animal biomass, supporting oceanic food webs. Despite their abundance and ecological importance, only five copepod genomes are publicly available, owing to a number of factors including large genome size, repetitiveness, GC-content, and small animal size. Here, we report the sixth representative copepod genome and the first genome and transcriptome from the calanoid copepod species *Acartia tonsa* Dana, which is among the most numerous mesozooplankton in boreal coastal and estuarine waters. The ecology, physiology and behavior of *A. tonsa* has been studied extensively. The genetic resources contributed in this work will allow researchers to link experimental results to molecular mechanisms. From PCRfree WGS and mRNA Illumina data, we assemble the largest copepod genome to date. We estimate *A. tonsa* has a total genome size of 2.5 Gb including repetitive elements we could not resolve. The non-repetitive fraction of the genome assembly is estimated to be 566Mb. Our DNA sequencing-based analyses suggest there is a 14-fold difference in genome size between the six members of Copepoda with available genomic information through NCBI. This finding complements nucleus staining genome size estimations, where 100-fold difference has been reported within 70 species. We briefly analyze the repeat structure in the existing copepod WGS datasets. The information presented here confirms the evolution of genome size in Copepoda and expands the scope for evolutionary inferences in Copepoda by providing several levels of genetic information from a key planktonic crustacean species.

## Introduction

Since the publication of the first version of the human genome sequence in 2001, more than 2000 eukaryotic genomes have been collected in the reference sequence database under NCBI [1]. The species with available genomic resources are predominately those which impact human health or are biomedically or agriculturally important. Genomic resources are available to a far lesser extent in species with ecological significance. Arthropoda is the most species rich phylum on Earth, and in marine environments, Copepoda is the most species rich subclass with more than 11,000 described species [2, 3], and the most abundant animal on Earth [4]. Yet, only five copepod genomes have hitherto been published. The five species are the calanoid *Eurytemora affinis* [5], the cyclopoid *Oithona nana* [6], the harpacticoid *Tigriopus californicus* [7] and *Caligus rogercresseyi* and *Lepeophtheirus salmonis*, which both belong to the order Siphonostomatoida and are important pests in salmon aquaculture [8].

*Acartia tonsa* is a marine, euryhaline calanoid copepod of about 1.5mm in adult length with a cosmopolitan neritic distribution, and in many ecosystems, it is the most numerous mesozooplankton species (Fig. 1A) [9]. It performs a vital function as it is a primary grazer on microalgae, and in turn is a main source of prey for the larvae of many fish species in estuarine, coastal and upwelling regions [10]. Further, *A. tonsa* is an emerging model organism, with research published in a diverse array of scientific fields such as ecology, physiology, ecotoxicology, and animal behavior [11–15]. *A. tonsa* is also an emerging live feed species in aquaculture, where it could trigger natural predation behavior and supply optimal nutrition for the larvae of fish species with economic importance or which are endangered in the wild [11, 16–18]. Despite *A. tonsa’s* multifaceted importance, partial versions of the mitochondrial cytochrome oxidase subunit I (COI) and the ribosomal 18S rRNA genes have been the only available genetic resources for *A. tonsa* until now [9, 19–21].

**Fig.1.**
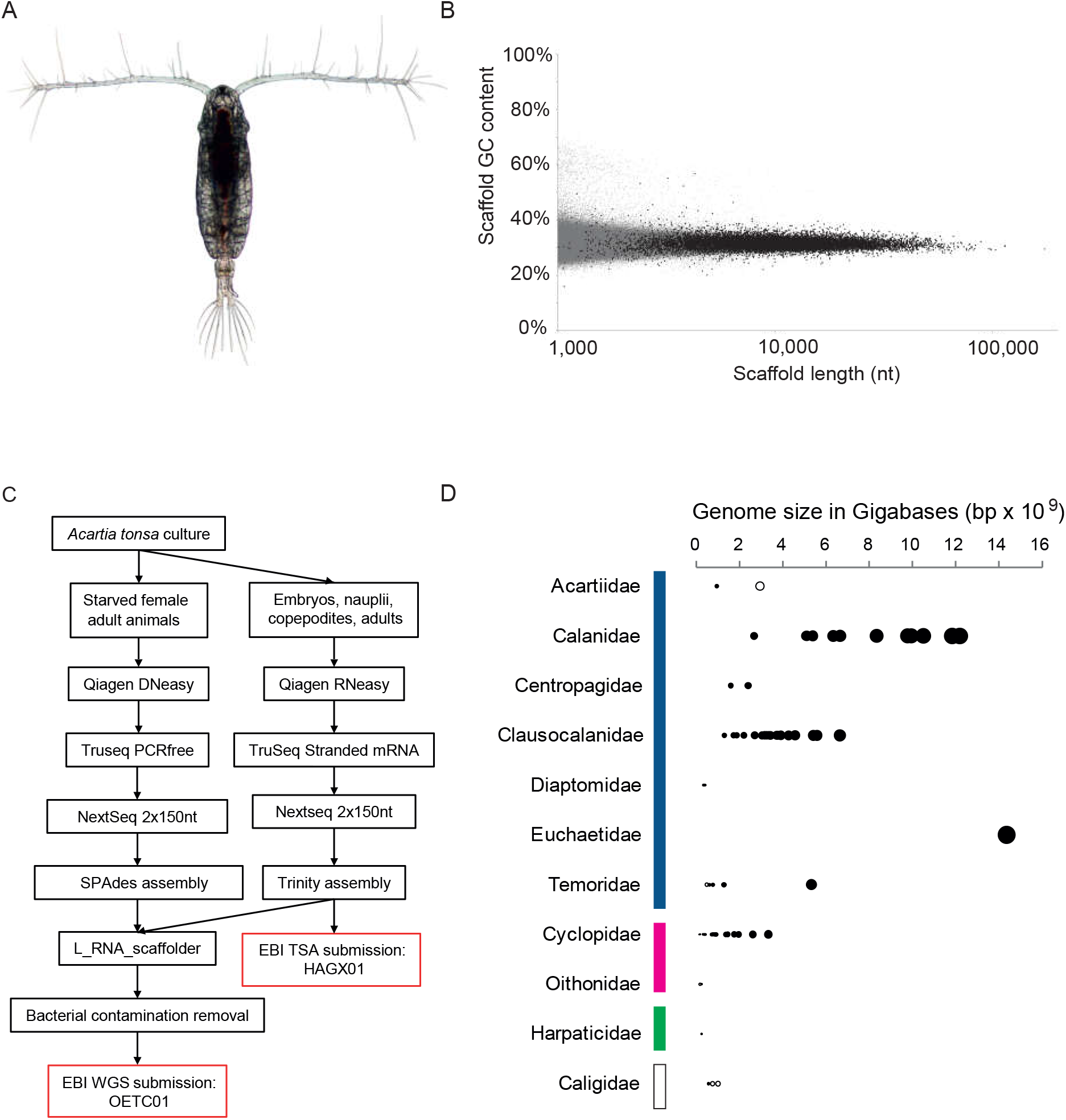
*A. tonsa* genome assembly. **A**, Female specimen of the DFU-ATI strain of *A. tonsa* used in this study. Photo by Minh Vu Thi Thuy. **B**, Length and GC-content of each scaffold in the Aton1.0 assembly. Black dots are scaffolds connected using mRNA information and grey dots are all other scaffolds. In total, 351,850 scaffolds are included in Aton1.0. The scaffolds are tightly distributed around 32% GC with lengths ranging from 1kb to 174kb. Most scaffolds of around or above 10kb have been scaffolded using mRNA information (Black dots). **C**, workflow for producing the Aton1.0 assembly from the DFU-ATI strain of *A. tonsa*. **D**, overview of reported genome sizes for the subclass Copepoda. The area of individual plot points is equal to the axis value. Black dots represent information from the Animal Genome Size Database based on nucleus staining (Gregory, T.R., 2018. http://www.genomesize.com) and the five open circles represent the genome sizes estimated from WGS data in this study. Within Copepoda, a 100-fold difference in genome size from the smallest Cyclopoid (pink bar, 0.14Gb) to the largest Calanoid (blue bar, 14Gb) can be seen. Within the order Calanoida, the genome sizes vary more than tenfold between the smallest Diaptomidae (0.95Gb) and the largest Calanidae (12Gb). Harpactidae species are marked with a green bar and Caligidae species with a white bar.

Copepod genomes are particularly difficult to assemble as the genomes are often large, have a very low GC content, around 30%, and because the animals are so small that a single animal rarely harbors a sufficient amount of genetic material for analysis [6, 22, 23]. This is compounded by the medical and agriculture focus of modern genome assembly pipelines, where diploidy, small genome size, abundant genetic material, and a GC content of ca 50% is assumed, required, or favored [24, 25].

Genome sizes of copepods species are highly variable. The genome assemblies available range 12-fold in size from 82 megabases (Mb) to 986 Mb (this study, Table 1), whereas the haploid genome sizes available from Feulgen staining of nuclei or flow cytometry range 100-fold between 140 Mb and 14 Gigabases (Gb) (Fig. 1D, Gregory, T.R., 2018. Animal Genome Size Database. http://www.genomesize.com). Within the order Calanoida, of which *A. tonsa* is a member, the reported haploid genome sizes vary 40-fold between 330 Mb and 14.4 Gb. For reference, the haploid human genome is ca 3.2 Gb [26]. The genome size of an animal is, however, not directly related to the assembly size, as the intronic and intergenic regions can be assembled to varying degrees which largely determine the assembly size [27]. Thus, we used all existing copepod Whole Genome Sequence (WGS) resources and our contributed *A. tonsa* genome assembly to determine the total genome size of copepods from the four orders of Calanoida, Cyclopoida, Siphonostomatoida, and Harpacticoida using the k-mer frequency based preQC tool. We further characterized the contributed genome of *A. tonsa* Dana by analyzing the content of mitochondrial marker genes. While few WGS datasets are available from Copepoda, transcriptome assemblies are much more common, with more than 20 datasets from 16 species available through the NCBI/EBI/DDBJ, likely owing to a relative ease of obtaining good quality transcriptomes compared to genomes. Our aim with this genome project was to contribute to the knowledge base of genome evolution in Copepoda and for the first time provide the research community with sufficient genomic and transcriptomic resources to embark on evolutionary, ecological, and physiological studies involving the important copepod species *A. tonsa*.

**Table1.**
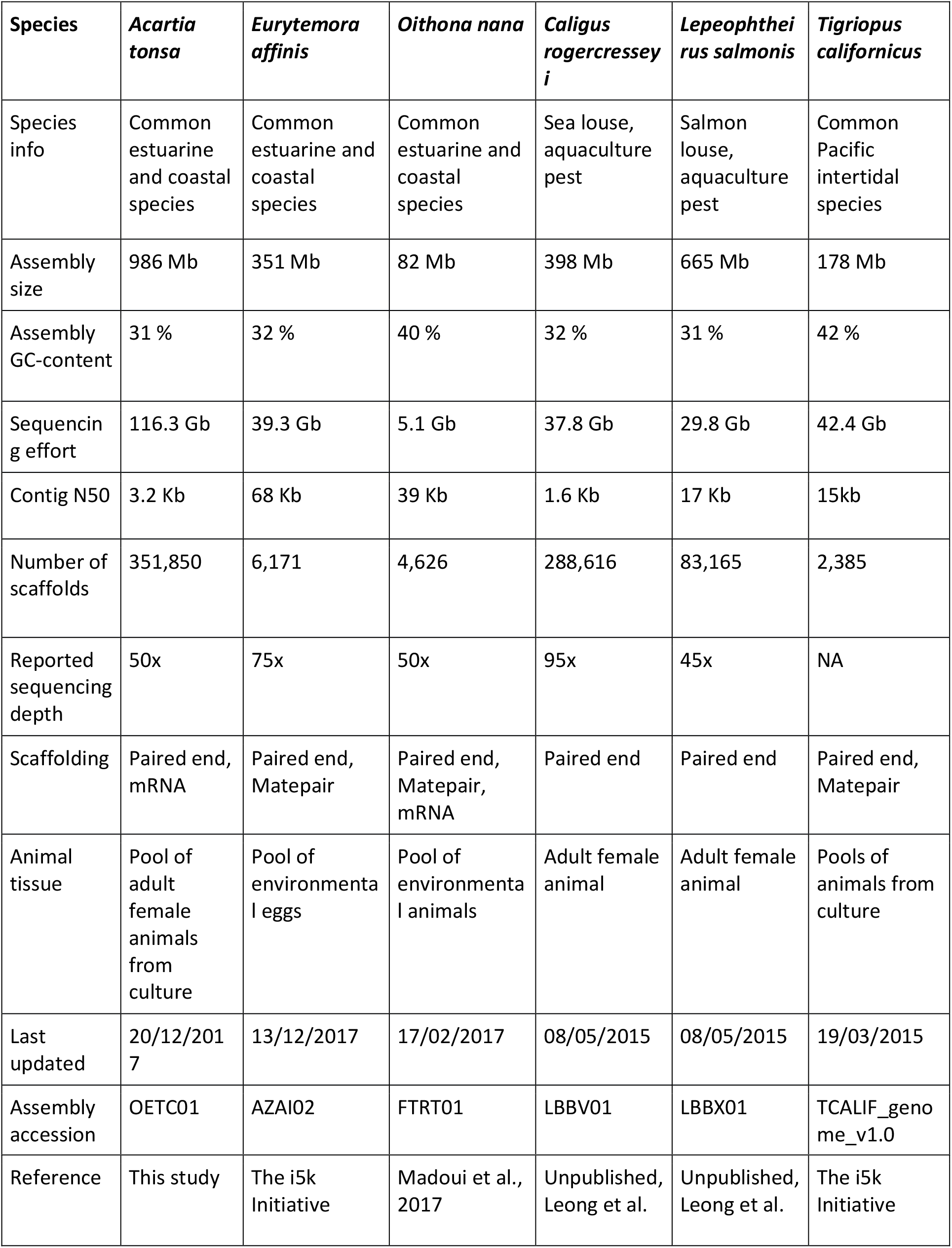
overview of existing genomic resources for Copepoda

## Results and discussion

### Sequencing and assembly metrics

The whole genome sequencing workflow for the *A. tonsa* genome (Aton1.0) and transcriptome (Fig. 1C) yielded a total of 356,383,864 Illumina reads from PCR-free libraries and 112,558,144 Illumina reads from three stranded mRNA libraries covering all life stages from embryos over nauplii and copepodites to adults. Further, PacBio and Mate Pair datasets were produced, but not used in the assembly process because the coverage was insufficient to successfully scaffold contigs (data not shown but available under the study accession PRJEB20069). The decision to not use the distance information libraries closely resembles the conclusions in a recent paper which reports that low coverage distance information does not improve assembly [28].

In total, more than 145,000,000,000 sequenced bases were used for the assemblies of *A. tonsa*, which is 3-5 times more raw data than any other copepod WGS study to date. The assembly of mRNA data yielded 118,709,440 bases in 61,149 transcripts and an additional 56,257 isoforms which are available at ENA (https://www.ebi.ac.uk/) under the accession HAGX01 (Fig. 1C). The SPAdes [29] assembly of PCRfree data was scaffolded with the transcriptome yielding a genome assembly of 989,163,677bp distributed in 351,850 scaffolds (Fig. 1B). The assembly is available at ENA (https://www.ebi.ac.uk/) under the name Aton1.0 and the accession OETC01. More than 20,000 contigs were joined using mRNA information, substantially adding to the contiguity of gene carrying scaffolds (Fig. 1B).

The GC content of the Aton1.0 assembly is 32% (Fig. 1B), substantially lower than many model species such as human or mouse but similar to the available Copepod genomes (Table 1). Because whole animals and embryos were used for nucleic acid extraction, bacterial contamination is expected to be present in the raw PCRfree data. The BLAST-based removal of scaffolds of bacterial origin eliminated 3,953 scaffolds, many of which had a substantially different sequence length and GC-content than other Aton1.0 scaffolds, further indicating bacterial origin (data not shown).

### Assembly completeness and content

To estimate the completeness of the Aton1.0 assembly, we used the BUSCO system of orthologous single copy genes and the arthropods database on predicted genes [30]. The Aton1.0 assembly carries 59.5% (634 of 1066) complete single copy BUSCO genes, 1.9% complete but duplicated genes (20 of 1066), 20.6% fragmented genes (220 of 1066), and 18.0% missing genes (192 of 1066) out of the 1,066 Arthropod gene models (Fig.2C). These numbers are not comparable with well-studied species such as *Drosophila melanogaster* (Fig. 2C, 99.0% complete, single copy genes) but are close to those of the other published copepods, though the sequencing effort for Aton1.0 is unprecedented (Fig. 2C and Table 1). The low number of duplicate BUSCO genes suggests that the Aton1.0 assembly is not populated by many variants of the same core genes, even though the biological material was obtained from a large number of animals. For the *A. tonsa* transcriptome, 91.4% of BUSCO genes are complete, and a further 7.9% fragmented, suggesting that this resource is very useful for scaffolding, gene modeling, and gene functional annotation. Genes annotation was done using MAKER2 [31] and both the mRNA transcriptome, ab initio gene prediction using Augustus, and related species gene models, but because the resulting gene set had a substantially lower BUSCO score than the *A. tonsa* assembly alone, we decided to not use it for further analysis (Supplementary material 1)

**Fig. 2.**
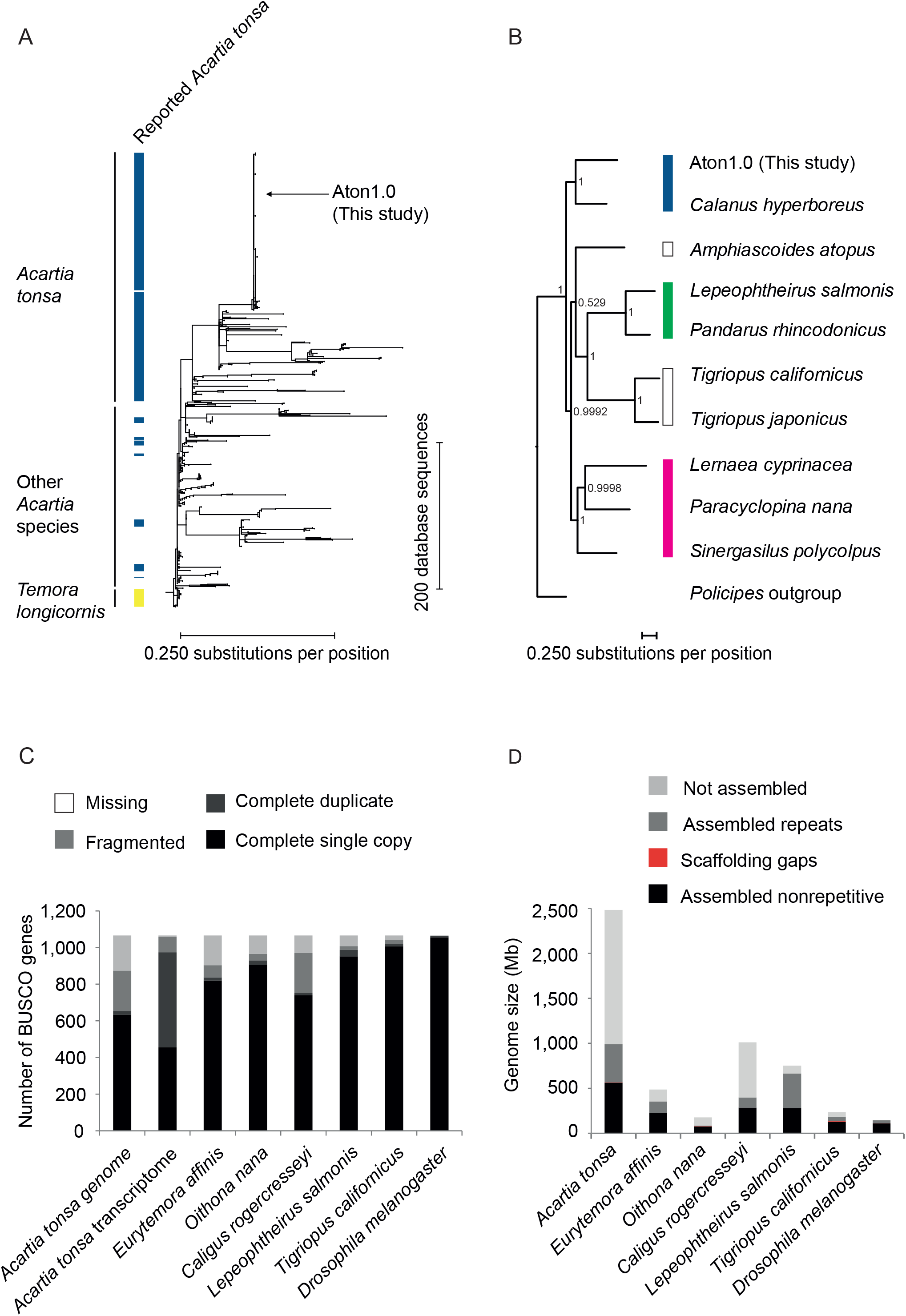
Placement and characterization of the Aton1.0 assembly. **A**, placement of Aton1.0 within the genus *Acartia*. The Aton1.0 COI gene groups within the most well studied North Atlantic clade of *A. tonsa* which is in line with the origin of the culture. **B**, placement of Aton1.0 within the subclass Copepoda. Phylognetic tree based on Bayesian analysis of a combined gene dataset. Nodal support is displayed as Bayesian posterior probability at each branch. The colored bars represent the orders Calanoida (blue), Harpacticoida (outline), Cyclopoida (cyan), and Siphonostomatoida (green). The branch separating the calanoid copepods from the other orders closely resemble recent phylogenetic analyses based primarily on different genes [33, 34] **C**, BUSCO core gene content of the genome assemblies of copepods and *D. melanogaster*. Between 2.4% and 18% of BUSCO genes are missing from assemblies (outline), 2.4% to 21% are fragmented (light grey), 1.2% to 3.2% exist in duplicate (dark grey). 59% to 94% of BUSCO genes are complete and single copy in the assemblies. For all metrics, the *A. tonsa* genome assembly performs worst, which is likely a result of the large genome size. The benchmarking species *D. melanogaster* has 99% complete single copy core genes. The mRNA transcriptome from all life stages of *A. tonsa* has 91% complete genes and additionally 8% fragmented genes, indicating that the resource is very powerful for identifying whole genes. **D**, Total genome sizes for all existing copepod WGS datasets and *D. melanogaster*. The unassembled genome fraction is depicted in light grey, the assembled repetitive genome fraction is in dark grey, the scaffolding gaps are in red and the non-repetitive assembled fraction in black. The Aton1.0 assembly represents a genome that is estimated to be three to 20 times larger than the other copepods for which WGS resources are available through NCBI. The fraction of assembled non-repetitive DNA is 22.7% (*A. tonsa*) to 53.8% (*T. californicus*) of the predicted total genome size, and only varies sevenfold from 75 Mb (*O. nana*) to 563 Mb (*A. tonsa*).

### Placement of Aton1.0 in *Acartia* and Copepoda

Mitochondrial genes and genomes are widely used for phylogenetic analysis in Copepoda because they often can resolve specimens to species [32]. Because one of the few sequences available from the DFU-ATI strain of *A. tonsa* is the mitochondrial cytochrome oxidase subunit I (COI) gene, we investigated the mitochondrial components in the Aton1.0 assembly. Three scaffolds were found to carry mitochondrial genes and they were annotated using MITOS2 [32] (Supplementary material 1). Within these three scaffolds, 15 out of 22 expected tRNA genes are present as well as 11 of 15 expected protein coding genes (table 2). The Aton1.0 COI gene was aligned to all 541 COI entries for the genus *Acartia* and a *de novo* phylogenetic tree was constructed based on a region shared between all database versions using 25 *Temora longicornis* (Copepoda, Calanoida) COI genes as outgroup (Fig. 2A). The Aton1.0 COI is 99.7% identical to many entries from the North Atlantic clade of *A. tonsa*, and most versions annotated as *A. tonsa* group together, confirming the placement of Aton1.0 within the most studied clade of *A. tonsa*, and the relatedness of database entries annotated as *A. tonsa*.

**Table 2.** overview of Aton1.0 mitochondrial resources

| Expected mitochondrial genes | Identified Aton1.0 mitochondrial genes | Used for phylogenetic placement |  | Expected mitochondrial tRNA | Identified Aton1.0 mitochondrial tRNA |
| --- | --- | --- | --- | --- | --- |
| ATP6 |  |  |  | trnA | A |
| ATP8 |  |  |  | trnC |  |
| COI1 | COI | x |  | trnD |  |
| COI2 | COI2 | x |  | trnE | trnE |
| COI3 | COI3 | x |  | trnF | trnF |
| CYTB | CYTB | x |  | trnG | trnG |
| ND1 | ND1 | x |  | trnH | trnH |
| ND2 |  |  |  | trnI |  |
| ND3 | ND3 | x |  | trnK | trnK |
| ND4 | ND4 | x |  | trnL1 | trnL1 |
| ND4L | ND4L |  |  | trnL2 | trnL2 |
| ND5 | ND5 | x |  | trnM |  |
| ND6 | ND6 |  |  | trnN | trnN |
| rRNA lsu | rRNA lsu |  |  | trnP | trnP |
| rRNA ssu |  |  |  | trnQ | trnQ |
|  |  |  |  | trnR |  |
|  |  |  |  | trnS | trnS1 |
|  |  |  |  | trnS | trnS2 |
|  |  |  |  | trnT |  |
|  |  |  |  | trnV |  |
|  |  |  |  | trnW | trnW |
|  |  |  |  | trnY | trnY |

Because nine complete circular copepod mitochondrial genomes are available, the Aton1.0 assembly can be placed within Copepoda using a multigene strategy. We chose eight complete genes for analysis, as these eight genes aligned across the database and our *A. tonsa* resource. Extreme mitochondrial DNA divergence has previously been reported even within the copepod species *T. californicus*, why it is not surprising that not all copepod mitochondrial genes align across orders [7]. The genes used for phylogenetic placement of *A. tonsa* can be found in Table 2. The result from the Bayesian phylogenetic analysis is presented in Fig. 2B. *A. tonsa* forms, together with *Calanus hyperboreus*, the clade of Calanoida (BPP:1), as a sister group to the clade of the remaining orders of Cyclopoida, Harpacticoida and Siphonostomatoida. *Paracyclopina nana, Lernaea cyprinacea* and *Sinergasilus polycolpus* forms the monophyletic clade of Cyclopoida (BPP:0.99). The paraphyletic clade of Harpacticoida and Siphonostomatoida is unsupported (BPP:0.53). Siphonostomatoida is nested as a monophyletic sister clade (BPP:1) to the clade of *Tigriopus japonicus* and *T. californicus* (BPP:1). The split between Calanoida and the other Copepod orders correlates closely with recent phylogenetic work on copepod orders where the calanoid species form the superorder Gymnoplea while the other copepods form the superorder Podoplea [33, 34]. The placement of the orders Cyclopoida, Harpacticoida, and Siphonostomatoida, however, is inconsistent in the recent papers, both of which are also inconsistent with our results, which suggests that Siphonostomatoida is nested inside Harpacticoida, with Cyclopoida as an outgroup (Fig. 2B). The cited studies use a larger number of species [33] or genes [34] for the analysis than the present work. We do not intent to challenge the validity of either, yet our result adds to the uncertainty of the placement of the copepod orders.

### Genome sizes and fractions

The total genome size of an animal including repetitive elements is not routinely deciphered from NGS data and reported along with the assembly. The preQC program from the SGA genome assembly pipeline uses k-mer frequencies to predict the total genome size and can be used to evaluate WGS data before assembly [35]. We tested preQC on the *D. melanogaster* genome. The estimated genome size of *D. melanogaster* is, within 3%, the same than the high-quality assembly genome length (Supplementary Material 1). This result permit us to use preQC on the five WGS genomes of copepods available at NCBI. For *A. tonsa*, the total genome size is estimated to be 2.48 Gb, slightly smaller than the size of the human genome (Fig. 2D). The almost 2.5 Gb genome size estimate of *A. tonsa* differs substantially from the other copepod genomes which are estimated to be 0.49Gb (*E. affinis*), 0.18 Gb (*O. nana*), 1.01 Gb (*C. rogercresseyi*), 0.75 Gb (*L. salmonis*), and 0.24 Gb (*T. californicus*). The complete preQC report for all species can be found in Supplementary Material 2. A 100-fold range in genome size has been reported in Copepoda based on nucleic staining (Fig. 1D), and the present study for the first time shows a large genome size range of 14-fold using NGS methods (Fig. 2D). Because the quality and quantity of input data can influence the result of k-mer counting based analysis such as the preQC genome size analysis, it is important to be aware that the genome size results are estimates, and that they could change with more input data, or data with a different error profile. A recent study used flow cytometry to estimate the genome size of four species of calanoid copepods, three of which also has Feulgen staining genome size estimates available [36]. The flow cytometry estimates were in all cases ca. half the size of the Feulgen staining estimates from the same species. This underlines the difficulty of copepod size estimations, and makes comparisons across methods difficult. Of the species analyzed in this study using an NGS method, *E. affinis, L. salmonis*, and *T. californicus* also have genome size estimates from a different method. For all, our estimates (0.49 Gb, 0.75 Gb, and 0.24 Gb, respectively) are close to the Feulgen staining estimates (0.62 Gb, 0.57 Gb, and 0.25 Gb, respectively) [37], Gregory, T.R., 2018. http://www.genomesize.com].

The large difference between the predicted genome sizes and the size of the genome assembly is hypothesized to be caused by both unassembled regions of the genome and the collapse of multiple repetitive regions to single scaffolds during assembly.

Because each assembly, scaffolding and gap filling approach yields different results, we determined the non-repetitive fraction of each available copepod genome by modeling and masking out repeats and analyzing total genome size, assembled repetitive sequence size, and non-repetitive sequence size. For *A. tonsa*, the non-repetitive fraction is 566Mb (Fig. 2D). This means that only 22.7 % of the *A. tonsa* genome is assembled and non-repetitive. This figure is substantially lower than for the other copepod species which have assembled non-repetitive fractions of 46.1% (224 Mb of 487 Mb), 42.4% (75 Mb of 177 Mb), 28.2% (285 Mb of 1011 Mb), 37.5% (282 Mb of 752 Mb), and 53.8% (127 Mb of 235Mb) for *E. affinis, O. nana, C. rogercresseyi, L. salmonis*, and *T. californicus*, respectively. This difference is possibly caused by the large genome size of *A. tonsa*, as larger genome size can be associated with increased amounts of repetitive DNA, while the amount of exon DNA remains stable [27]. Figure 3 shows the amount of classified and unclassified repeats in the copepod genomes. Characteristically, the large majority of repeats in all copepod genomes cannot be classified by the RepeatMasker program [38] using RepeatModeler [39] output combined with the Repbase_arthropoda database (downloaded 2017-06-02). For *Drosophila*, most repeats are classified as long terminal repeats (LTR, 40 % of repeats) or long interspersed repeats (long interspersed nuclear element, LINE 17% of repeats, Fig. 3), whereas only 25% of identified repeats could not be classified. Likely, the *D. melanogaster* repeat classification is much better than the copepod repeat classification because *D. melanogaster* is a model species which specifically has been included in the RepBase repository. The WGS assembly of *T. californicus* is among the most contiguous copepod genome assemblies, and a larger fraction of repeats from this species can be classified by Repeatmasker than from the other copepod species (Fig.3). Still, even for *T. californicus*, almost 60 % of the repeats could not be classified. For A. tonsa, 95 % of the identified repeats could not be classified, which is the highest rate of any of the analyzed copepods (Fig. 3). The largest amount of classified repeats in the *A. tonsa* assembly is simple repeats, which make up 17Mb or 4 % of the identified repeats. This is equivalent to <1% of the total genome length. It is important to consider the large unassembled fraction of most copepod genomes when analyzing repeat structure, as the sequence absent from assemblies are very likely to be repetitive DNA and as the missing genome fraction constitute up to 60 % of the total genome length.

**Fig. 3.**
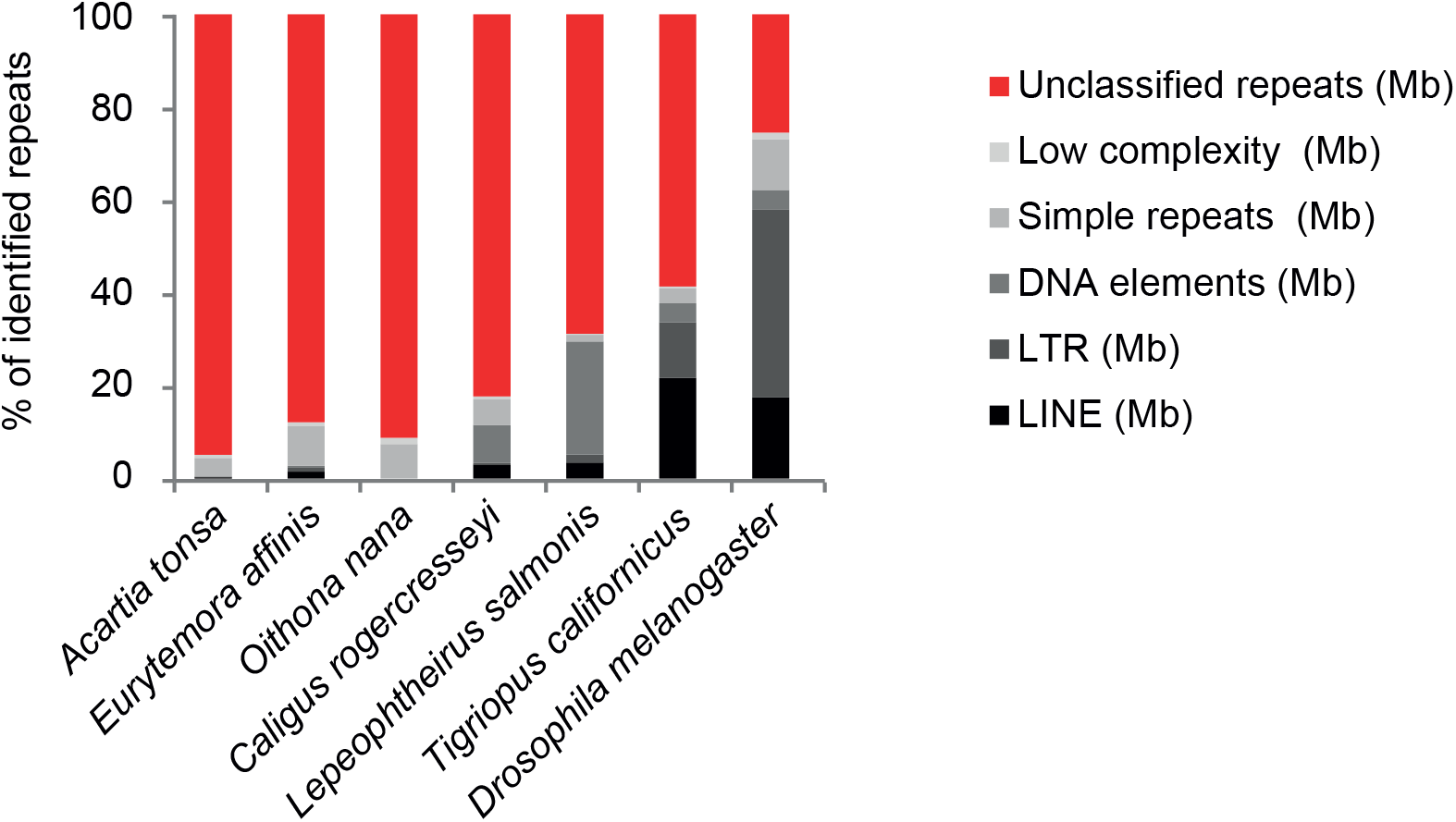
Classification of repeats in copepod WGS assemblies using RepeatModeler and RepeatMasker. While more than 70% of identified repeats can be classified in the model species *Drosophila*, only between 5 % and 20 % of identified repeats from copepod genomes was classified. The unassembled genome fractions described in Fig 2D and the large amount of unclassified repeats in copepods together illustrates how limited the current knowledge on this important animal group is.

## Conclusions

Here, we present the first transcriptome and genome assembly of the ecologically important copepod species *Acartia tonsa* Dana. 82 % of the BUSCO core genes are present in the genome assembly, including 2 % duplicated and 21 % fragmented genes. In the transcriptome assembly 99 % of BUSCO genes could be found, including 8 % fragmented genes. We further document the placement of the contributed genome within Copepoda and the genus *Acartia* to the North Atlantic clade and estimate the genome size of *A. tonsa* to almost 2.5 Gb and compare to the other available copepod genomic resources where we find a 14-fold difference in estimated genome size. This is the first documentation of the range of genome size within Copepoda using DNA sequencing methods. Our resources are likely valuable to researchers in many scientific fields and can assist others to consider genome size when planning genome sequencing projects by understanding causal bases for the difference between the genome size and the assembly size of animal genomes.

## Materials and methods

### Culture and animal husbandry

The *A. tonsa* culture strain DFU-ATI was used for all nucleic acid extractions. DFU-ATI has been in continuous culture without restocking since it was obtained off the coast of Helsingør in the Øresund strait in Denmark in 1981. Behavioral, ecological, physiological, and molecular aspects of the biology of *A. tonsa* strain DFU-ATI have been described in several publications [11–14, 40–43]. The continuous *A. tonsa* culture fed the microalga *Rhodomonas salina* in excess according to [44] was kept in 70L plastic buckets in a stable 17 °C environment in the dark. The culture was kept in 0.2 μm filtered water collected from the sea floor in Kattegat, near the site where the culture originated. The salinity was stable at 32 ±1 ppt. Eggs and debris were collected from the bottom of the culture daily.

Animals were sorted by size by sequential filtering: adults were caught on a 250 μm filter, copepodites and nauplii were caught on a 125 μm filter and eggs were caught on a 70 μm filter. Animals were thoroughly rinsed with 0.2 μm filtered seawater which was removed prior to nucleic acid extraction. For the PCRfree libraries, individual adult animals were picked with a Pasteur pipette and placed in sterile, 0.2 μm filtered seawater which was removed prior to nucleic acid extraction. Tissues for RNA extraction were placed in at least 5 volumes of RNAlater 24h prior to extraction.

### Nucleic acid extraction, library construction, and sequencing

DNA was extracted using the DNeasy mini Blood and Tissue kit from Qiagen (Hilden, Germany) according to the manufacturer’s protocol with the following modifications: sample tissue was ground manually with a pestle in a 1.5 ml Eppendorf tube for at least two minutes and incubated with proteinase K for 4 hours with periodic mixing.

RNA was extracted using the RNeasy mini Blood and Tissue kit from Qiagen (Hilden, Germany) according to protocol with the following modifications: sample tissue was kept on ice and ground in a 1.5 ml Eppendorf tube using a pestle and electric motor for two minutes and incubated with proteinase K for 4 hours with periodic mixing. The embryo RNA sample consisted of eggs from a few and up to 50 hours old to ensure that all stages in *A. tonsa* embryogenesis were present [45].

The PCRfree libraries were constructed using DNA from adult female animals with the Truseq PCR free kit (Illumina, San Diego, CA, USA) according to the manufacturer’s protocol using 1 μg DNA as input. Shearing of DNA in the PCR free protocol was done on a Covaris E210 (Woburn, MA, USA) with miniTUBE with the following settings: Intensity: 3, duty cycle: 5%, cycles/burst: 200, Treatment time: 80. The libraries were sequenced on an Illumina NextSeq 500 with 2×150 bp PE high kits. The three transcriptome libraries covering all life stages of *A. tonsa* were built using the Illumina TruSeq stranded mRNA kit with half volumes according to [46] immediately after RNA extraction. No DNase step was used as it would have negative impact on long fragments in the libraries. For each library, 1 μg of total RNA was used as input. RNA libraries were sequenced on an Illumina Nextseq 500 using a single 2×150 bp PE high kit. All sequencing libraries were analyzed on a Bioanalyzer with DNA 7500 Assay chip (Santa Clara, CA, USA) and the molarity of cluster forming fragments was analyzed using the KAPA Universal qPCR Master Mix (KK4824, KAPA Biosystems, Wilmington, MA, USA). Nucleic acid concentration was measured using the Qubit system (Thermo Fisher Scientific, Waltham, MA, USA).

An overview of the libraries was constructed, and more details on indexes, insert sizes, biological materials, and SRA accession numbers can be found in Supplementary Material 1.

### Data handling, assembly, scaffolding, and analysis

Basic statistics and data handling was done in a UNIX environment using Biopieces (Hansen, MA, www.biopieces.org, unpublished). mRNA data from eggs, nauplii, copepodites, and adults was pooled and assembled using the Trinity pipeline (v. 2.5.1) with default parameters and the built-in version of trimmomatic to quality trim the reads prior to assembly and to remove adapters [47, 48]. Data from PCRfree libraries was pooled and used for assembly with SPAdes assembler [29] (v. 3.9.0, k-mer size 77), using paired end information for scaffolding. The SPAdes genome assembly was further scaffolded with the assembled mRNA transcriptome using L_RNA_scaffolder [49] with BLAT v. 36×2 [50]. Transcripts shorter than 500 nt were not used for scaffolding. Scaffolds smaller than 1,000 bp were discarded. After testing several bacterial contamination removal strategies, we decided on a BLAST based method on scaffold level sequences as all sequences with obvious bacterial characteristics (high GC% and 100Kb-1Mb in length) were removed and few likely copepod sequences were removed (data not shown). Briefly, scaffolds were masked using RepeatMasker with repeats from RepeatModeler and the Arthropoda and ancestral (shared) repeats from repbase v. 22.05 (downloaded 2017-06-02) [38, 39]. The masked scaffolds were BLAST-searched against the refseq database of representative prokaryotes (downloaded 2017-03-23) using the built-in BLAST in CLCgenomics 9.0 (e-value ≤10^−6^) [51] and sequences with a longest hit longer than 500 bp were removed from the assembly. Raw reads and assemblies for the four published copepod reference genomes were downloaded from NCBI using the following accession numbers: *E. affinis* (assembly: AZAI02, raw reads: SRX387234-7), *O. nana* (assembly: FTRT01, raw reads ERX1858579-83), *C. rogercresseyi* (assembly: LBBV01, raw reads: SRX976492), *L. salmonis* (assembly: LBBX01, raw reads SRX976783), and *T. californicus* (assembly TCALIF_genome_v1.0, raw reads SRX469409 and SRX469410). Genome size was estimated using the preQC tool from the SGA assembler [35] on reads cleaned using AdaptorRemoval [52] with the switches: --trimns–trimqualities. Repetitive sequence fractions in genome assemblies were identified using RepeatMasker on the Arthropoda and ancestral (shared) repeats from repbase v. 22.05 (downloaded 2017-06-02) merged with output from RepeatModeler run with standard parameters [38, 39].

The nine complete copepod mitochondrial genomes used to find mitochondrial scaffolds in Aton1.0 were downloaded from the Organelles section of the NCBI genomes browser (https://www.ncbi.nlm.nih.gov/genome/browse#!/organelles/copepoda) in April 2018. Their accession numbers can be found in Supplementary Material 1. BLAST search of the nine existing mitochondrial genomes against the Aton1.0 assembly was done in CLC Genomics workbench v. 10.1.1 using standard parameters and yielded three scaffolds with mitochondrial genes. The scaffolds carrying mitochondrial DNA were analyzed using the MITOS2 web interface with RefSeq63 Metazoa reference and table 5 invertebrate genetic code [32].

COI genes from the genus *Acartia* along with 25 *Temora longicornis* COI genes were downloaded from the NCBI nucleotide collection (https://www.ncbi.nlm.nih.gov/nuccore/?term=Acartia%20COI) in April 2018. The accession numbers of the 544+25 sequences can be found in Supplementary Material 1. Multiple alignment of Aton1.0 and database COI, trimming of sequence ends, realigning and *de novo* Neighbor Joining phylogenetic tree construction with 100 bootstraps were all performed in CLC genomics workbench version 10.1.1 using default parameters. Analysis of genome and transcriptome completeness was done using the Universal Single Copy Orthologs BUSCO (v2.0) with the arthropoda_odb9 lineage dataset and *ab initio* gene prediction using Augustus (v.3.2.3), in all cases with the fly training set, and the switch ‘geno’ for the genomes and ‘tran’ for the transcriptome [30, 53].

To place Aton1.0 within Copepoda, the eight genes COX1, COX2, COX3, CYTB, ND1, ND3, ND4, and ND5 were extracted from the MITOS annotation of Aton1.0 scaffolds and from the nine complete copepod mitochondria downloaded from NCBI (Supplementary Material 1). The genes were aligned individually using the MAFFT online platform [54], using the algorithm Q-INS-I iterative refinement method [55]. Individual genes were then concatenated using Sequence Matrix [56].

The concatenated dataset of the mitochondrial genes was analyzed using Bayesian method (BA). The analysis was performed using MrBayes version 3.2.6 [57] available on CIPRES Gateway. To identify the best substitution model for molecular evolution a Model Test was run on CLC Genomics Workbench 10.1.1 (https://www.qiagenbioinformatics.com/) on each individual gene prior to analyses, using Akaike information criterion (AIC) for COX2, COX3, CYTB, ND1, ND3 and corrected Akaike information criterion (AICc) for COX1, ND4, ND5. The models selected for each gene included a General time reversible (GTR) model of sequence evolution [58] with gamma distribution and a proportion of invariable sites (GTR + I + Γ) for all genes. The dataset was run with two independent analyses using four chains (three heated and one cold). Number of generations was set to 30 million, sampling every 1,000 generations. Burn-in was set to 10 million generations.

## Acknowlegdements

We thank Anna la Cour for excellent technical assistance (ORCHID ID: 0000-0002-6990-9891), and Marlene Danner Dalgaard (ORCID ID: 0000-0002-4036-6408) for advice and assistance with DNA shearing. This study was supported by the Villum Foundation project AMPHICOP 8960.

## Data accessibility

Raw DNA sequencing data, the genome assembly, and the transcriptome assembly are available under the project PRJEB20069. The genome assembly prefix for Aton1.0 is OETC01 and the transcriptome prefix is HAGX01. All further data is available in Supplementary Material 1 or upon request.

## Author contributions

TSJ, BHW, LHH, JHS designed research, TSJ performed research, BP, SP, JHS, PDB, HCP contributed analytical tools, TSJ, BP, SP, JHS, HCP analyzed data, TSJ, BP, SP, JHS, HCP, PDB, LHH, BHW wrote the paper.

