## Supplementary figures and images for "The genome and mRNA transcriptome of the cosmopolitan calanoid copepod *Acartia tonsa* Dana improve the understanding of copepod genome size evolution"

### Supplementary material 2

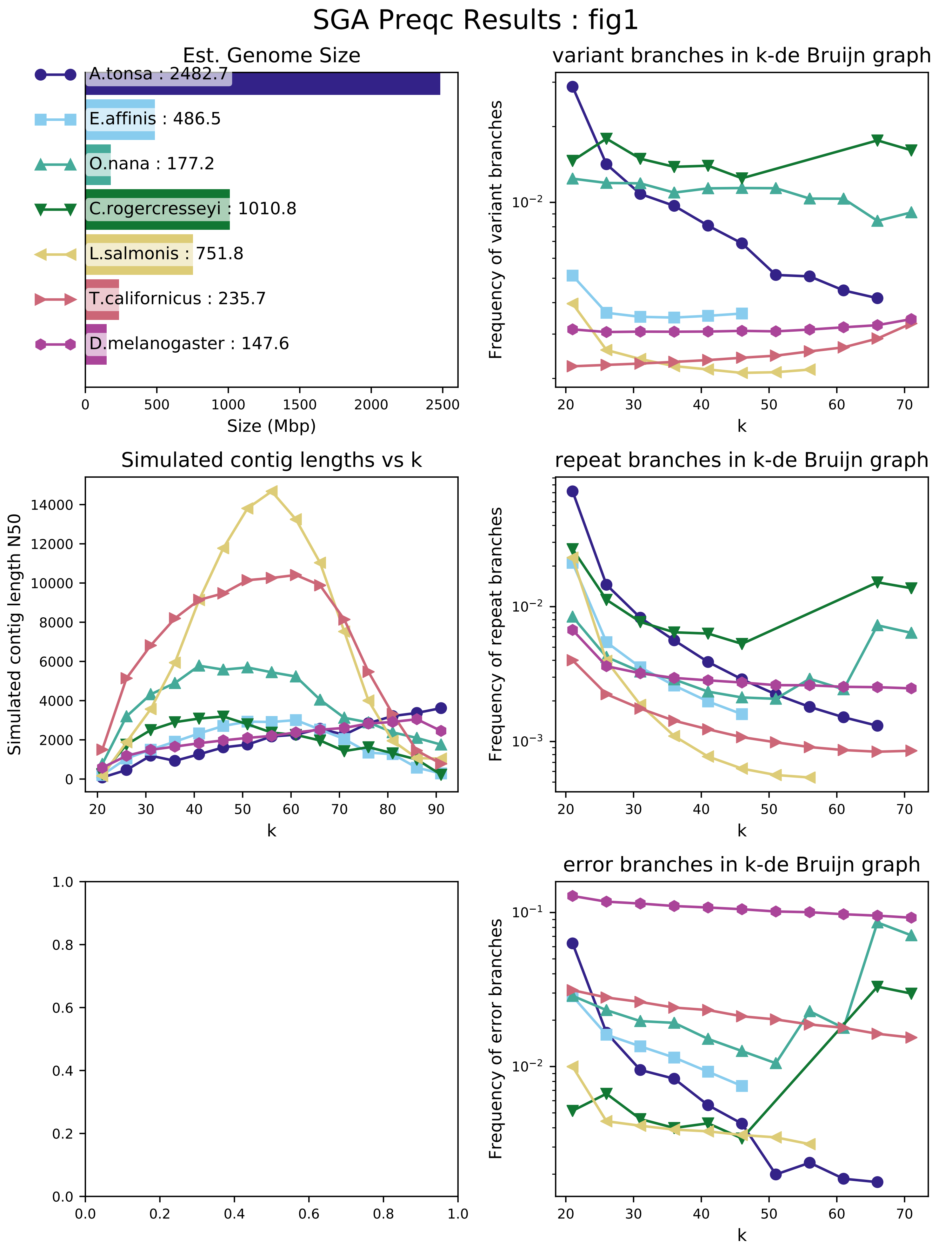

#### SGA Preqc Results : fig2

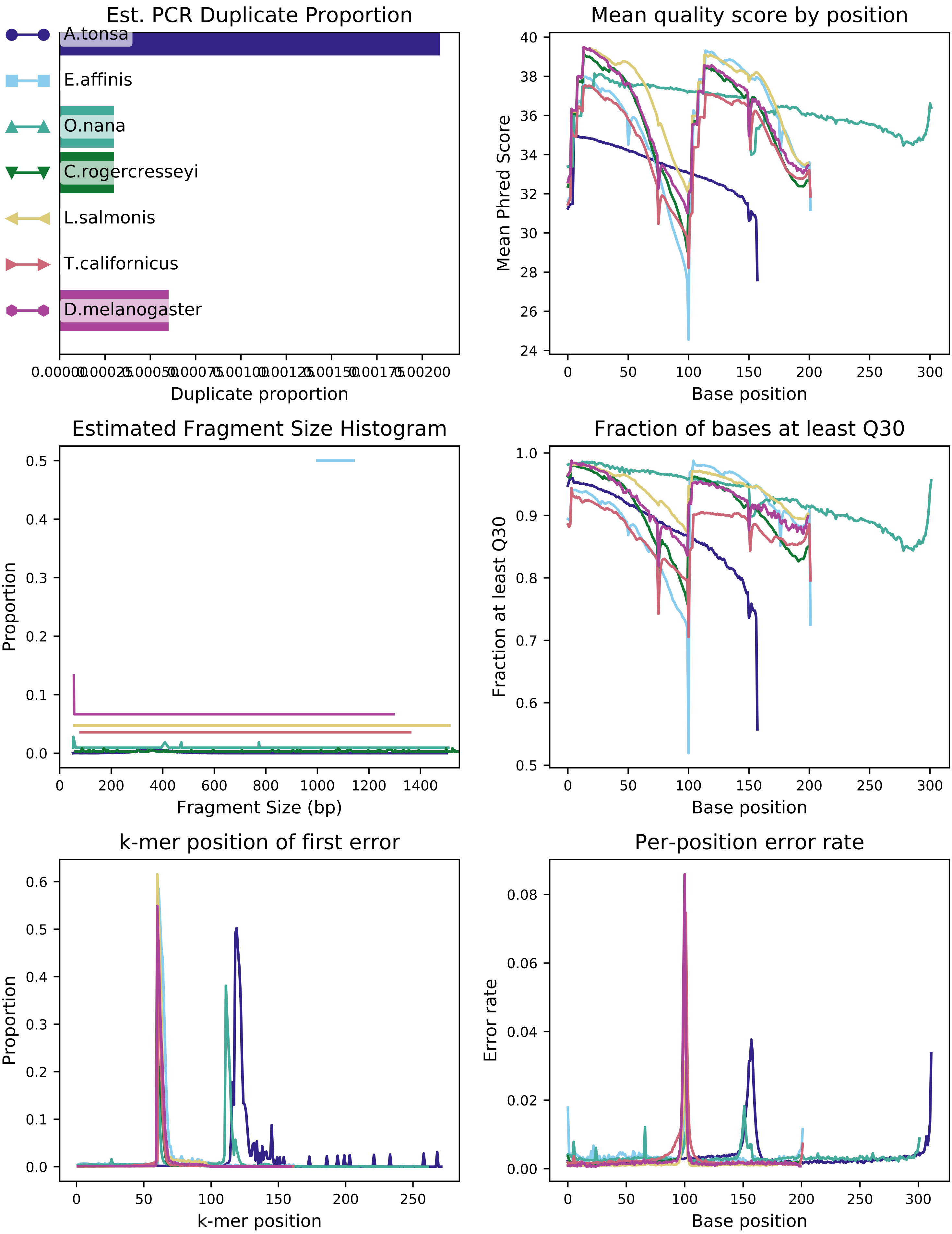

#### SGA Preqc Results : fig3

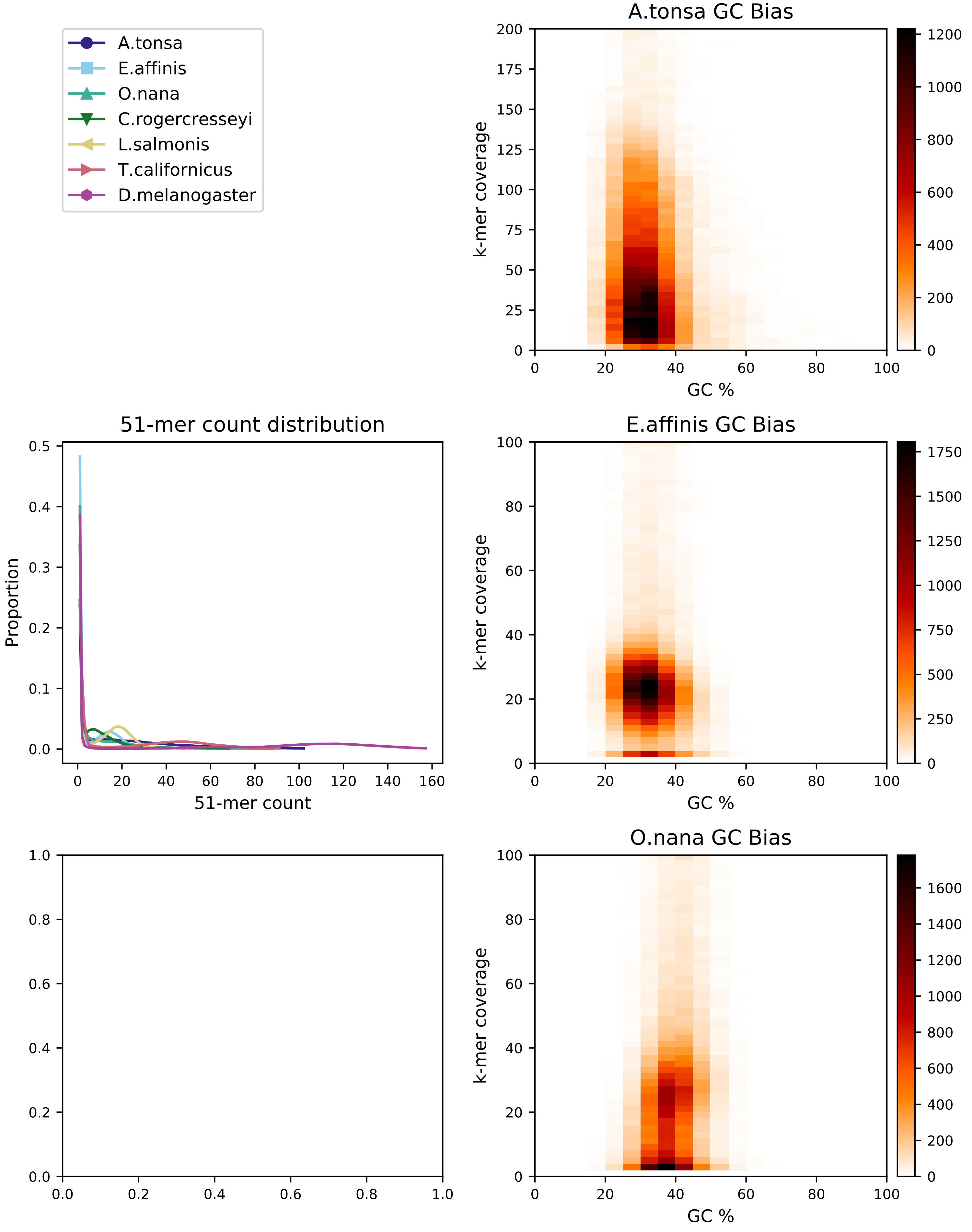

#### SGA Preqc Results : fig4

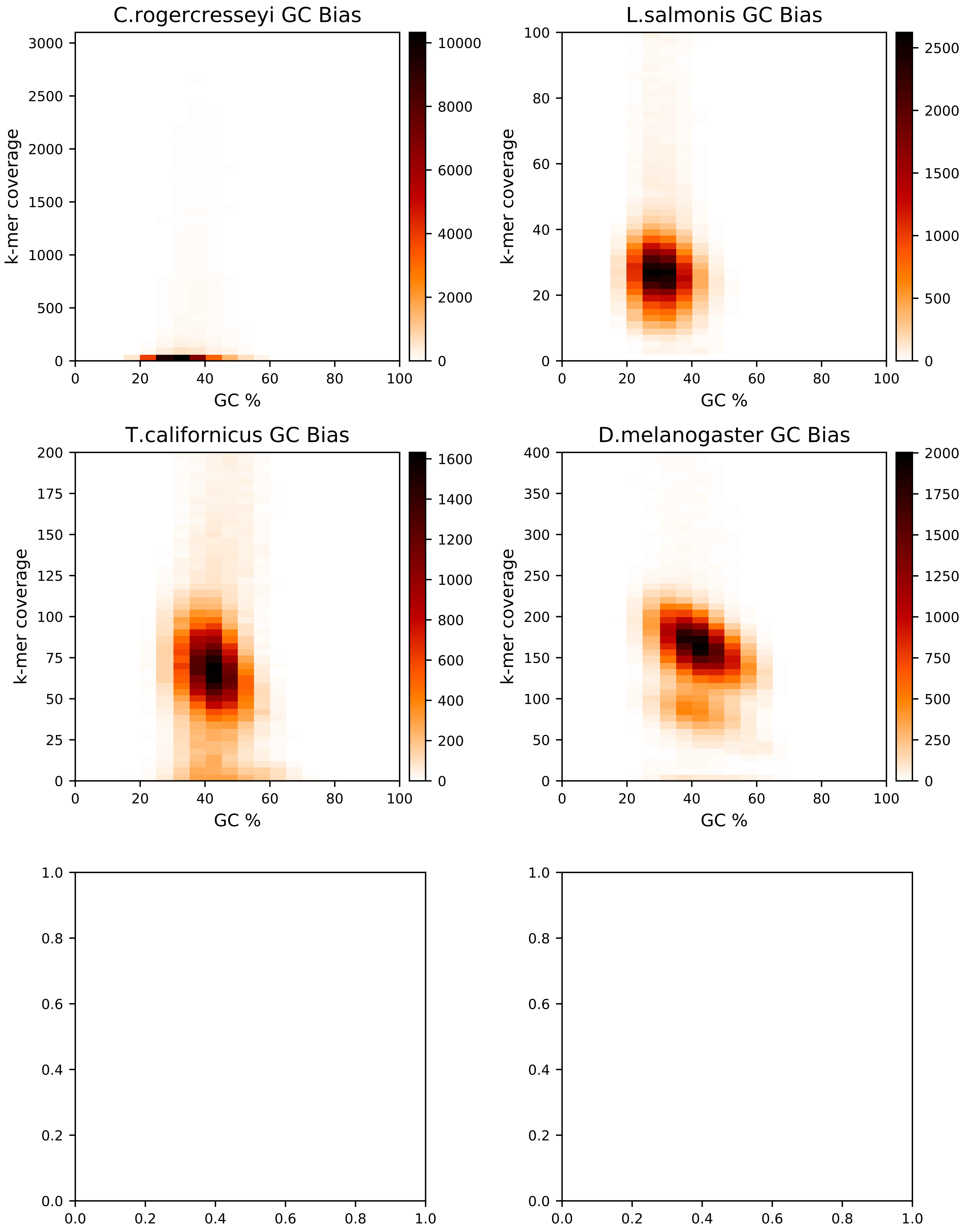
